# Subgenome dominance in an interspecific hybrid, synthetic allopolyploid, and a 140 year old naturally established neo-allopolyploid monkeyflower

**DOI:** 10.1101/094797

**Authors:** Patrick P. Edgar, Ronald Smith, Michael R. McKain, Arielle M. Cooley, Mario Vallejo-Marin, Yaowu Yuan, Adam J. Bewick, Lexiang Ji, Adrian E. Platts, Megan J. Bowman, Kevin L. Childs, Robert J Schmitz, Gregory D. Smith, J. Chris Pires, Joshua R. Puzey

**Author notes:** These authors contributed equally (Patrick P. Edgar, Joshua R. Puzey).

## Abstract

The importance and applications of polyploidy have long been recognized, from shaping the evolutionary success of flowering plants to improving agricultural productivity. Recent studies have shown that one of the parental subgenomes in ancient polyploids is generally more dominant - having both retained more genes and being more highly expressed - a phenomenon termed subgenome dominance. How quickly one subgenome dominates within a newly formed polyploid, if immediate or after millions of years, and the genomic features that determine which genome dominates remain poorly understood. To investigate the rate of subgenome dominance emergence, we examined gene expression, gene methylation, and transposable element (TE) methylation in a natural less than 140 year old allopolyploid (*Mimulus peregrinus*), a resynthesized interspecies triploid hybrid (*M. robertsii*), a resynthesized allopolyploid (*M. peregrinus*), and diploid progenitors (*M. guttatus* and *M. luteus*). We show that subgenome expression dominance occurs instantly following the hybridization of two divergent genomes and that subgenome expression dominance significantly increases over generations. Additionally, CHH methylation levels are significantly reduced in regions near genes and within transposons in the first generation hybrid, intermediate in the resynthesized allopolyploid, and are repatterned differently between the dominant and submissive subgenomes in the natural allopolyploid. Our analyses reveal that the subgenome differences in levels of TE methylation mirror the increase in expression bias observed over the generations following the hybridization. These findings not only provide important insights into genomic and epigenomic shock that occurs following hybridization and polyploid events, but may also contribute to uncovering the mechanistic basis of heterosis and subgenomic dominance.

## 1 Introduction

Whole genome duplications (WGD) have been an important recurrent process throughout the evolutionary history of eukaryotes^1–3^, including having contributed to the origin of novel traits and shifts in net diversification rates^4–7^. WGDs are especially widespread across flowering plants^8–10^, with both deep WGD events (all extant angiosperms share at least two events^11^) and a plethora of more recent events including those unique to our model system *Mimulus^12^.* Polyploids, species that have three or more complete sets of genomes, are grouped into two main categories: autopolyploids (WGD that occurred within a species) and allopolyploids (WGD coupled with a interspecific hybridization)^13^. Previous studies indicate that allopolyploids are more likely to persist and become ecologically established - a fact that has partially been attributed to heterosis due to transgressive gene expression and fixed heterozygosity^3, 7, 14, 15^. Newly formed allopolyploids face the unique challenge of organizing two genomes (i.e. subgenomes), each contributed by different parental species, that have independently evolved in separate contexts, which now exist within a single nucleus^16^. Hybridization and allopolyploidization may disrupt both genetic and epigenomic processes resulting in altered DNA methylation patterns^17–23^, changes in gene expression^24–28^ and transposable element (TE) reactivation^29^ - commonly referred to as genomic shock^30^. These genome wide changes are associated with novel phenotypic variation in newly formed allopolyploids^31, 32^, which likely contributed to the survival and ultimate success of polyploids^33^.

One observation that may be linked to the long-term success of allopolyploids is that homeologous genes (homologous genes encoded on different parental subgenomes) are often expressed at non-equal levels, with genome-wide expression abundance patterns being highly skewed towards a subgenome. Examples of plants with evidence for subgenome specific expression include Maize^34^, *Brassica*^35^, cotton^26^, wheat^36^, *Tragopogon*^24^, *Spartina*^25^, and *Arabidopsis*^37^. Additionally, it has been shown that the less expressed subgenome tends to be more highly fractionated (i.e. accumulate more deletions), a pattern thought to be due to relaxed selective constraints. Collectively these phenomena are referred to as ‘subgenome dominance’^38^. For example, in *Brassica rapa*, a three-way battle ensued following a whole genome triplication event that occurred over ten million years ago resulting in a single dominant subgenome emerging and two highly fractionated subgenomes^39^. The most highly fractionated subgenome lost more than double the total number of genes than the least fractionated, dominant subgenome.

It remains largely unknown how one subgenome becomes more highly expressed, with respect to either whole genome patterns or specific genes. Another unanswered question is, on what time scale (i.e. how quickly) does subgenome dominance become established? Subgenome dominance in newly formed hybrids and allopolyploids could have substantial implications for our understanding of plant hybridization in both ecological and agricultural contexts. In addition, a mechanistic understanding of these phenomena is fundamental to better understanding the long-term evolutionary advantages of WGDs. One hint at a mechanism may be that gene expression can be impacted by the proximity to and methylation status of nearby TEs^40^. Prompted by the finding that the density of methylated TEs is negatively correlated with gene expression magnitude, Freeling et al.^41^ hypothesized that the relationship between TE repression and the expression of neighboring genes might explain patterns of observed subgenome dominance. The degree of methylation repatterning and reestablishment genome-wide, specifically nearby genes, following hybridization and/or WGD is largely unknown. Here we tested this hypothesis by assessing (i) the overall rate that subgenome expression dominance is established following hybridization and WGD, (ii) genome-wide methylation repatterning following hybridization and WGD, and (iii) the influence of methylation repatterning on biased expression of parental subgenomes.

Most polyploid systems are hindered by at least one of two major difficulties: (1) lack of genomic resources for extant parental progenitors (if parents are known) or (2) the inability to confidently partition the polyploid genome to each of the parental subgenomes. Here we used the recently formed natural allopolyploid, *Mimulus peregrinus*, to overcome these hurdles. *M. peregrinus* (6x) is derived from the hybridization of *M. luteus* (4x) and *M. guttatus* (2x), which produced a sterile triploid intermediate *M. x robertsii* (3x) that underwent a subsequent WGD to regain fertility^12, 42^ (Fig 1). Importantly, *M. luteus* (native to Chile) and *M. guttatus* (native to Western North America) only recently came into contact following a documented introduction into the UK in the early 1800’s^12^. Thus, we have a narrow time window for the formation of *M. peregrinus.* Moreover, the natural allopolyploid (*M. peregrinus*) still exists with its introduced parents in the UK, which allows us to recreate hybrids and synthetic allopolyploids in lab. Furthermore, the *M. guttatus* genome was recently published^43^ and we complement this with a new genome assembly for *M. luteus* (see Results). These resources and the unique natural history of *M. peregrinus* have provided an unprecedented opportunity to properly investigate subgenome expression dominance and its relation to DNA methylation and TE density differences.

**Figure 1:**
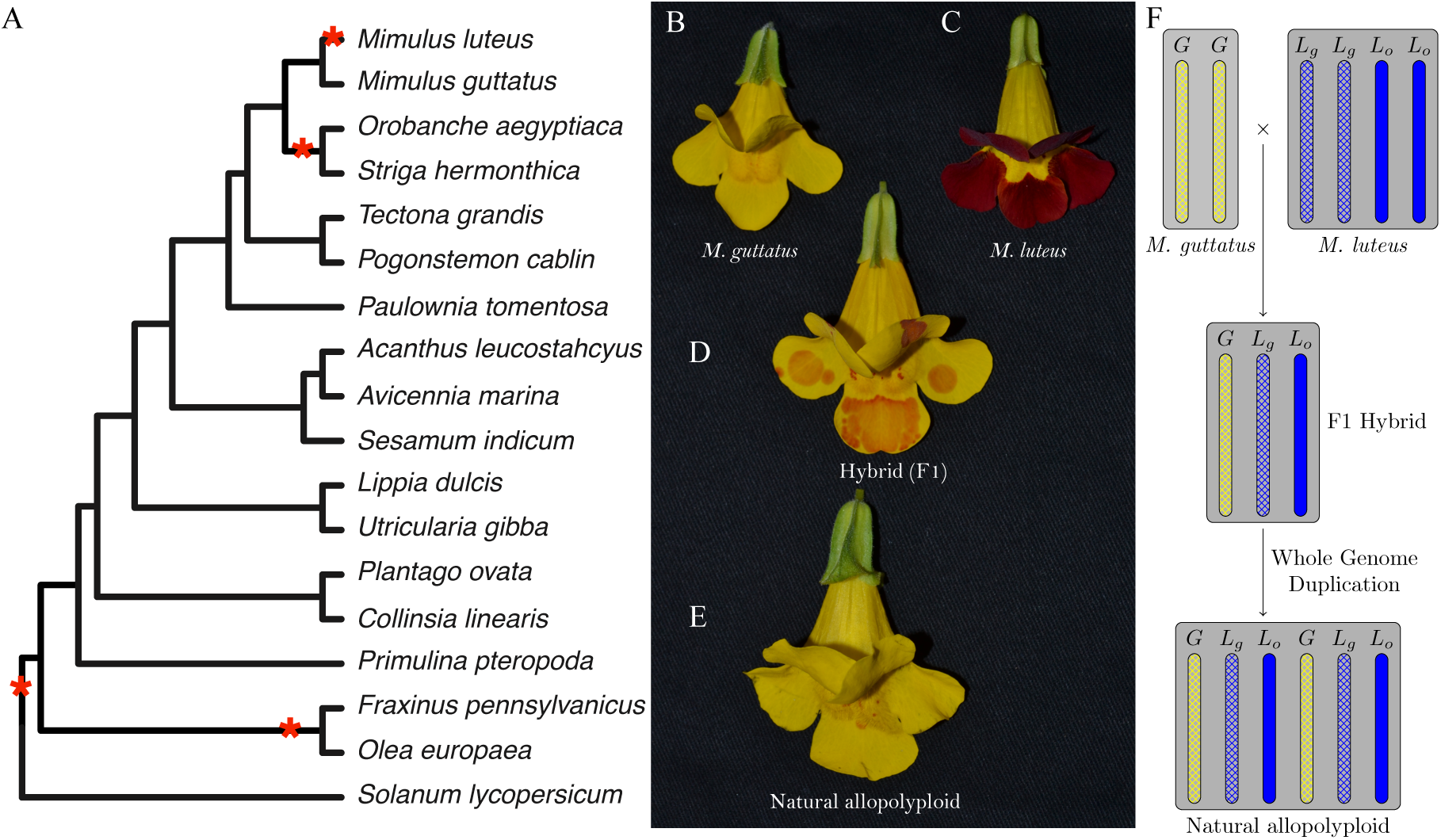
Whole genome duplications in *Mimulus* and related species. (A) Tree showing locations of whole genome duplications (asterisks) on Lamiales phylogeny. (B) Mimulus species used in this study. *Mimulus guttatus* [2x] hybridized with (C) *Mimulus luteus* [4x] to produce a sterile triploid (D) *Mimulus robertsii* [3x] which underwent a subsequent whole genome duplication giving rise to fertile natural allopolyploid (E) *Mimulus peregrinus* [6x]. (F) Graphic showing chromosome complement of individuals (B-E) in middle panel. The allotetraploid *M. luteus* has two distinct subgenomes; *L_g_* represents the *M. guttatus-like* subgenome and *L_o_* represents the ‘other’ subgenome.

## 2 Results

### *Mimulus luteus* genome assembly

Here we present the draft genome of *M. luteus* with a total genome size estimate of 640-680 megabases (Mb) based on flow cytometry and kmer spectrum analysis (SI Appendix, Text S1.1)^42^. The assembly contains 6,439 scaffolds spanning 410Mb with an N50 of 283kb, representing roughly 60 percent of the genome, with gene content analyses supporting the recovery of nearly the entire gene space (SI Appendix, Text S1.1). A total of 46,855 protein coding genes were annotated in *M. luteus* genome, which is nearly double the number of protein coding genes (26,718) previously annotated in the *M. guttatus* genome (430 Mb estimated; 300 Mb assembled)^43^. This difference in gene content supports a tetraploid event (WGD) unique to *M. luteus*, previously reported based on both genome size and base chromosome number differences between these species^44^. We re-annotated the *M. guttatus* genome with identical methods used for *M. luteus*, reducing the total number to 25,465 protein coding genes (SI Appendix, Text S1.1). The re-annotation of this genome permits us to make proper genomic and transcriptomic comparisons by removing artifacts that arise due to differences in genome annotation pipelines. A total of 319,944 and 451,448 TEs were annotated in the *M. guttatus* and *M. luteus* genomes, respectively (SI Appendix, Text S1.6). We combined these two genomes to represent *M. peregrinus.*

### History of WGD in *Mimulus*

A shared ancient whole genome duplication was detected in both genomes, termed *Mimulus*-alpha, with a mean Ks of 0.92 and phylogenetically placed at the most recent common ancestor of *Mimulus* (Phrymaceae) and nearly all other Lamiales families (Fig 1) (SI Appendix, Text S1.2, Figs. S1, S2, and S3). A recent mean-date estimate for that phylogenetic node is 71 million years before present^45^. Our taxon sampling did not include the earliest diverging lineage (family Plocospermataceae) in Lamiales^45^. The Mimulus-alpha event is shared by all other surveyed Lamiales families. A total of 757 unique shared duplications from Mimulus-alpha were identified in all surveyed taxa (SI Appendix, Text S1.2). Two additional duplication events detected across Lamiales were not shared with Phrymaceae; First, a WGD event shared by *Orobanche* (Orobanchaceae) and *Striga* (Oleaceae) supported by 1739 shared duplicate pairs. Second, a WGD event shared by *Olea* and *Fraxinus* supported by 3308 shared duplicate pairs (SI Appendix, Text S1.2).

*Mimulus luteus* experienced an additional whole genome duplication event not shared with *M. guttatus.* As a result, for every *M. guttatus* gene, *M. luteus* typically has two corresponding homeologs. We sought to determine whether the two *M. luteus* homeologs were more similar to each other or whether one *M. luteus* homeolog was consistently more similar to the *M. guttatus* ortholog. This analysis was prompted by the finding of Mukherjee and Vickery (1962) that *M. luteus* is an allopolyploid formed by the hybridization of a *M. guttatus-like* individual and a non-*M. guttatus* individual^44^. Within the *M. luteus* genome, we identified 2200 high-confidence duplicate pairs that coalesce to the common ancestor of *M. luteus* and *M. guttatus.* Next, we compared these homeologs to each other and to their respective *M. guttatus* homeolog. Through this analysis we determine that in 1853 of the 2200 cases, one of the *M. luteus* homeologs was more similar to its *M. guttatus* homeolog than it was to its *M. luteus* paralog. Below, the *M. luteus* homeologs more similar to the *M. guttatus* homeolog is denoted as *‘M.* guttatus-like’, while the remaining *M. luteus* paralog will be referred to as ‘other’. Of the remaining 347 *M. luteus* paralogs, 341 pairs formed a clade sister to the *M. guttatus* homolog. The remaining six *M. luteus* pairs were unclear. Thus, using homeologous *M. luteus* genes and their orthologous *M. guttatus* gene as identified through a whole genome synteny (SI Appendix, Text S1.2), we find that one of the *M. luteus* homeologs is significantly more similar to *M. guttatus* than it is to the other *M. luteus* homeolog.

### Investigating the establishment of subgenome dominance

RNA-seq datasets for calyx, stem, and petals were generated for the F1 hybrid *M. x robertsii (M. guttatus* x *M. luteus*), resynthesized allopolyploid *M. peregrinus*, and naturally derived allopolyploid *M. peregrinus* and parental taxa, *M. luteus* and *M. guttatus* (SI Appendix, Text S1.3). These data were used to measure homeolog specific gene expression following the *M. luteus* WGD event as well as in a contemporary hybrid and neo-allopolyploids. All RNA samples were collected within a narrow time range to control for major diurnal rhythmic expression differences. In hybrids and allopolyploids, subgenome-specific (parental) SNPs were identified and were used to measure homeolog specific gene expression (SI Appendix, Text S1.3). For the analysis of our RNA-seq data, we developed a likelihood ratio test (LRT) involving three nested hypotheses to identify cases of homeolog expression bias that do not involve tissue specific expression differences (SI Appendix, Text S1.5). The null hypothesis is that both homeologs are expressed at equal levels (ratio of homeolog-1 to homeolog-2 equals 1 for all three tissues). The first alternate hypothesis is that homeologs are expressed at different levels, but similar ratios, across all three tissue types. The second alternate hypothesis is that homeologs are expressed at different levels and at different ratios across all three tissues.

Using the expression data and the nested hypotheses we test for: (1) Expression bias and subgenome dominance following the *M. luteus* specific WGD in *M. luteus*, hybrid *M. guttatus x M. luteus*, and *M. peregrinus* (both natural and resynthesized allopolyploid) and (2) Expression bias and subgenome dominance of the *M. guttatus* or *M. luteus* homeologs in the hybrid and allopolyploid *M. peregrinus.* Importantly, the first comparison will allow us to understand long term patterns of expression bias in *Mimulus* (the *M. luteus* WGD is a relatively ancient event) whereas the second comparison will test for expression bias and subgenome dominance in a newly formed hybrid and allopolyploid *Mimulus*.

### Expression bias of homeologs from *Mimulus luteus* specific WGD event

Based on the finding that one of the *M. luteus* subgenomes is significantly more like *M. guttatus*, we sought to compare expression of homeologs within *M. luteus* to each other, thereby addressing the question of whether the *M. guttatus-like* homeolog or ‘other’ homeolog is more highly expressed within *M. luteus, M. x robertsii* (F1 hybrid), or *M. peregrinus* (both a resynthesized and natural allopolyploid). Using the likelihood ratio test mentioned above we identified cases of homeolog expression bias that did not involve tissue specific expression differences. In each case over 1100 homeolog pairs were tested; the test was only applied when both homeologs were expressed in all three tissues. For each homeolog pair, we quantified expression bias as 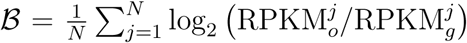 where the subscripts g and o denote the *M. guttatus-like* and ‘other’ subgenome respectively, and *j* is an index over the *N* tissues. An expression bias of 𝓑 = −2 indicates a 4x expression bias towards the *M. guttatus-like* homeolog, while an expression bias of 𝓑 = 3 is 8x towards the ‘other’ homeolog.

The histograms in Figure 2 summarize the measured expression bias of homeologs resulting from the *M. luteus*-specific WGD event in *M. luteus*, F1 hybrid, synthetic allopolyploid, and natural allopolyploid. For *M. luteus* (top panel) the grey histogram shows the distribution of expression bias for all testable homeolog pairs indicating a slight average bias towards the *M. guttatus-like* subgenome (𝓑̅ = −0.08, *N_Lg_* = 364 > 329 = *N_Lo_*). Using the likelihood ratio test developed in this paper, we found that over half of the homeolog pairs in *M. luteus* (*N_Lg_* + *N_Lo_* = 693, about 51% of the total *N*) were biased towards one of the subgenomes with no tissue specific expression differences (hypothesis one, see Methods). Of these, a small majority of homeologs (*N_Lg_* = 364, about 54% of *N*), were dominately expressed from the *M. guttatus*-like subgenome. In contrast to *M. luteus*, in the hybrid, synthetic allopolyploid, and natural allopolyploid, the ‘other’ subgenome is slightly dominant in both number and average (see panels two through four where about 26%, 30%, and 29% of homeologs are dominately expressed from the ‘other’ subgenome and 𝓑̅ =0.06, 0.03, and 0.04, respectively).

**Figure 2:**
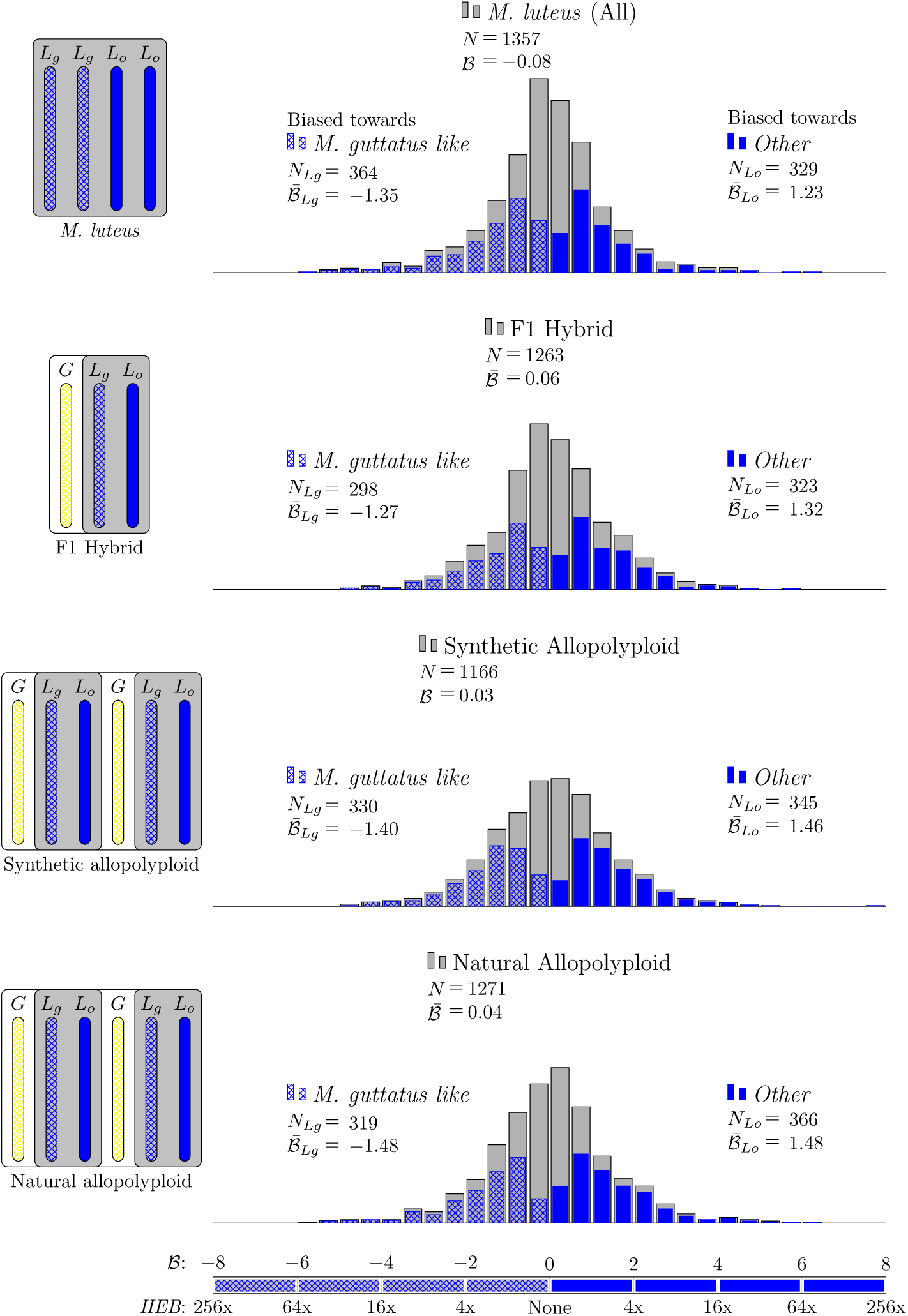
Expression bias of homeologs resulting from *M. luteus* specific WGD event in *M. luteus*, F1 hybrid, synthetic allopolyploid, and natural allopolyploid. Gray histograms show distribution of expression bias (𝓑) for all testable homeolog pairs. Testable homeolog pairs (*N*) are those which could clearly be identified as homologous and had at least 1 read in each tissue sampled. Homeolog pairs significantly biased towards the *M. guttatus-like* homeolog are crosshatched, while pairs significantly biased towards the ‘other’ homeolog are shown in solid blue. Across all three hybrid individuals (F1, synthetic, and natural allopolyploid) the ‘other’ subgenome dominates the *M. guttatus-like* subgenome either by the number of homeologs biased towards it (*N_Lo_* > *N_Lg_*) or on average, 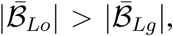 where 𝓑̅*_Lo_* and 𝓑̅*_Lg_* are averages over all homeolog pairs that were biased towards *L_o_* or *L_g_*, respectively.

### Expression bias of *M. guttatus* and *M. luteus* homeologs in the hybrid, resynthesized allopolyploid, and naturally occurring neo-allopolyploid *Mimulus*

To test for expression bias that arises instantaneously following the merger of two genomes, we compared homeologs in the hybrid and neo-allopolyploids, which contain both a *M. guttatus* and *M. luteus* subgenome. We asked two questions. First, when considering *M. luteus* expression as the sum of its two homeologs (*L_o_* + *L_g_*), do we see dominance of one subgenome (Fig. 3)? Second, when we consider the *M. luteus* homeologs separately (*L_o_* or *L_g_*) and compare these to their *M. guttatus* homeolog (*G*), do we see do we see dominance of one subgenome? The answer to the first question may indicate which parental subgenome overall is contributing the most gene product while the second question will shed light whether one single subgenome is more dominant. For each homeolog pair, we quantified expression bias as 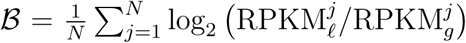 where the subscripts *ℓ* and *g* denote the *M. luteus* and *M. guttatus* subgenome respectively, and *j* is an index over the *N* tissues.

**Figure 3:**
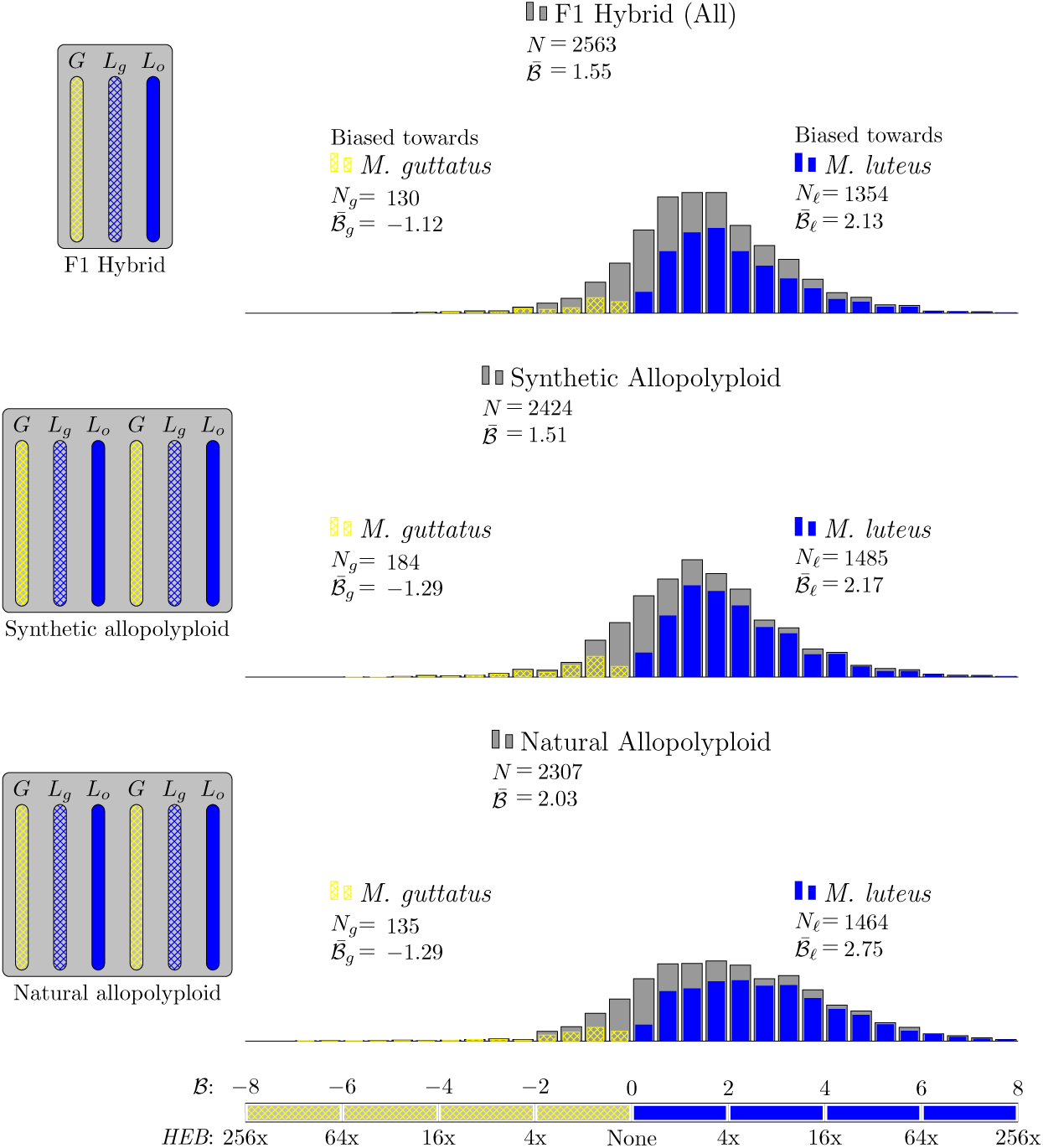
Homeolog expression bias in hybrid and allopolyploids, comparing the *M. guttatus* homeolog to the sum of its pair of *M. luteus* homeologs. Gray histograms show distribution of expression bias (𝓑) for all testable homeolog pairs. Only genes which had a clear 2 to 1 (*M. luteus* to *M. guttatus*) homology were considered. Homeolog pairs significantly biased towards the *M. guttatus* homeolog are shown in yellow, while pairs significantly biased towards the *M. lutues* homeolog are shown in blue. Across all three hybrid individuals (F1, synthetic, and natural allopolyploid) the pair of *M. luteus* homeologs, when added together, dominates the *M. guttatus* homeolog (i.e., *N_ℓ_* > *N_g_* and 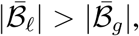 where 𝓑̅*_l_* and 𝓑̅*_g_* are averages over all homeolog pairs.)

The histograms in Figure 3 summarize the measured expression bias in hybrid and allopolyloids, comparing the expression of the *M. guttatus* homeolog to the sum of the expression of its pair of *M. luteus* homeologs. On average, there is considerable bias towards the *M. luteus* subgenome with 𝓑̅ = 1.55, 1.51, and 2.03 in the first generation hybrid, synthetic allopolyploid, and natural allopolyploid, respectively. Using the LRT (at significance level *α* = 0.01) a total of 1484 (58%), 1669 (69%), and 1599 (69%) homeolog pairs were found to be significantly biased. Of these biased pairs, the *M. luteus* homeolog was dominate in the vast majority of cases (*N_ℓ_* = 1354 (91%), 1669 (89%), 1599 (92%), respectively).

The histograms in Figure 4 summarize the measured expression bias in hybrid and allopolyloids, comparing the expression of the *M. guttatus* homeolog to expression of each of its *M. luteus* homeologs separately. When considering the *M. luteus* homeologs separately, expression is still considerably biased on average towards *M. luteus* with 𝓑̅ = 0.48, 0.46, and 0.97, in the first generation hybrid, resynthesized allopolyploid, and natural allopolyploid, respectively (Fig. 4). Next, using the LRT, across all comparisons, 52%, 63%, and 66% homeologs were significantly biased. Of these biased homeolog pairs, the *M. luteus* homeolog was the dominantly expressed homeolog in 68%, 64%, and 73% of the comparisons (Fig. 4). Additionally, among biased homeologs, the average bias towards the *M. luteus* subgenome is greater than the average bias towards the *M. guttatus* subgenome 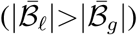 in all cases (Fig. 4). It is also worth noting that the degree of bias (as measured by 𝓑 and fraction of biased homeologs) increased from the first generation hybrid, to the resynthesized allopolyploid, and to the natural allopolyploid.

**Figure 4:**
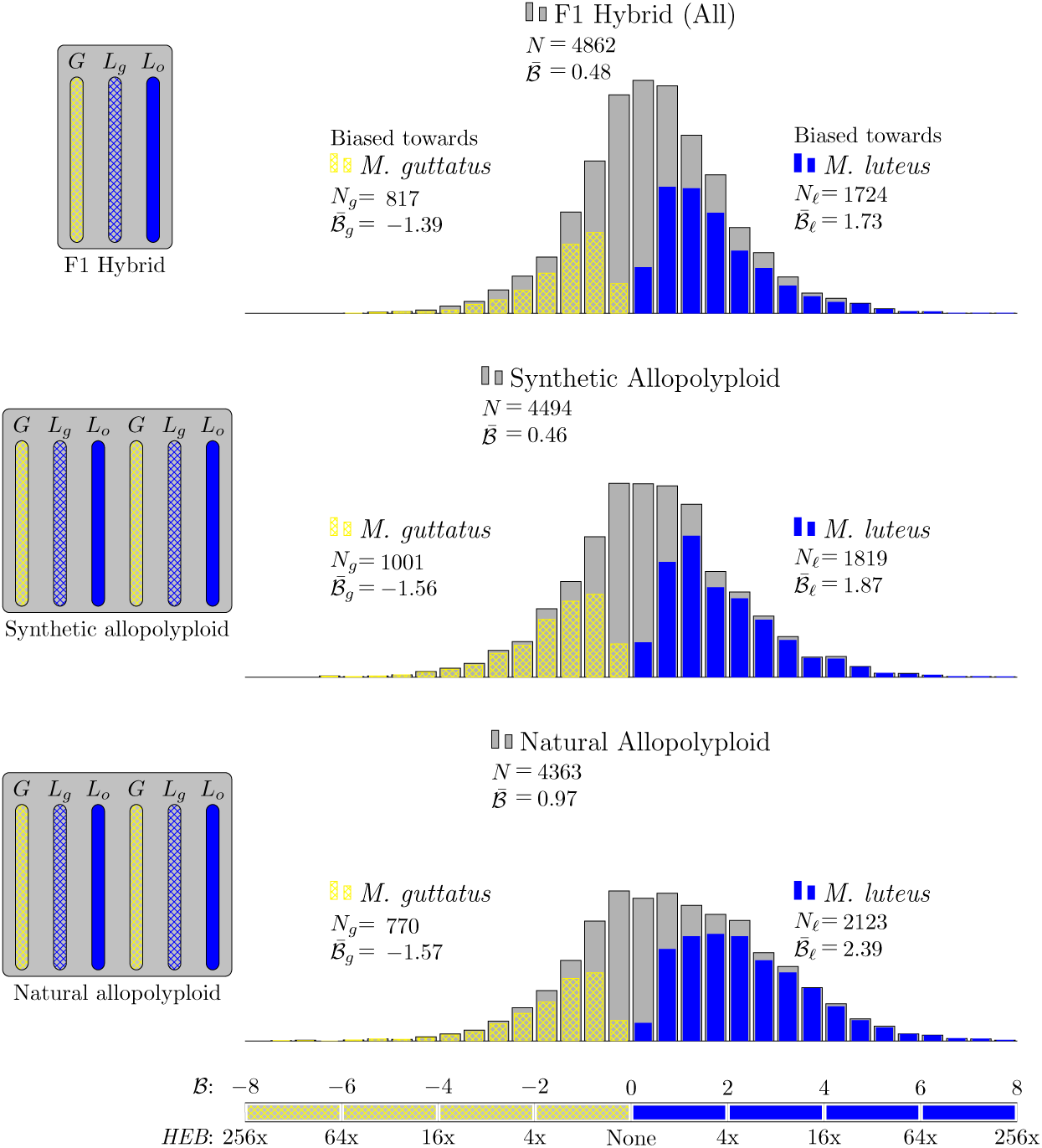
Homeolog expression bias in hybrid and allopolyploids, comparing the *M. guttatus* homeolog to each of its *M. luteus* homeologs separately. Gray histograms show distribution of expression bias (𝓑) for all testable homeolog pairs. Homeolog pairs significantly biased towards the *M. guttatus* homeolog are shown in yellow, while pairs significantly biased towards the *M. lutues* homeolog are shown in blue. Across all three hybrid individuals (F1, synthetic, and natural allopolyploid) the *M. luteus* homeolog dominates the *M. guttatus* homeolog (i.e., *N_ℓ_* > *N_g_* and 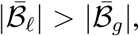 where 𝓑̅*_l_* and 𝓑̅*_g_* are averages over all homeolog pairs.)

#### Expression bias in three separate hybrid lineages

While it is clear that the *M. luteus* homoelogs are dominantly expressed in the hybrid an allopolyploid lineages, we sought to determine whether the same homeologs were repeatedly biased across independent hybrid and allopolyploids. A Venn diagram reveals that homeologs biased in one individual are far more likely to be biased in the other two lineages than would be expected by random chance (Fig. 5). Moreover, measured levels of individual homeolog expression bias are correlated across all three lineages (Fig. 5). Interestingly, levels of expression bias 𝓑 in the first generation hybrid and resynthesized allopolyploid are much more correlated with each other (*r*^2^ = 0.57) than either sample is with the natural allopolyploid (*r*^2^ = 0.25 and 0.32, Fig. 5).

**Figure 5:**
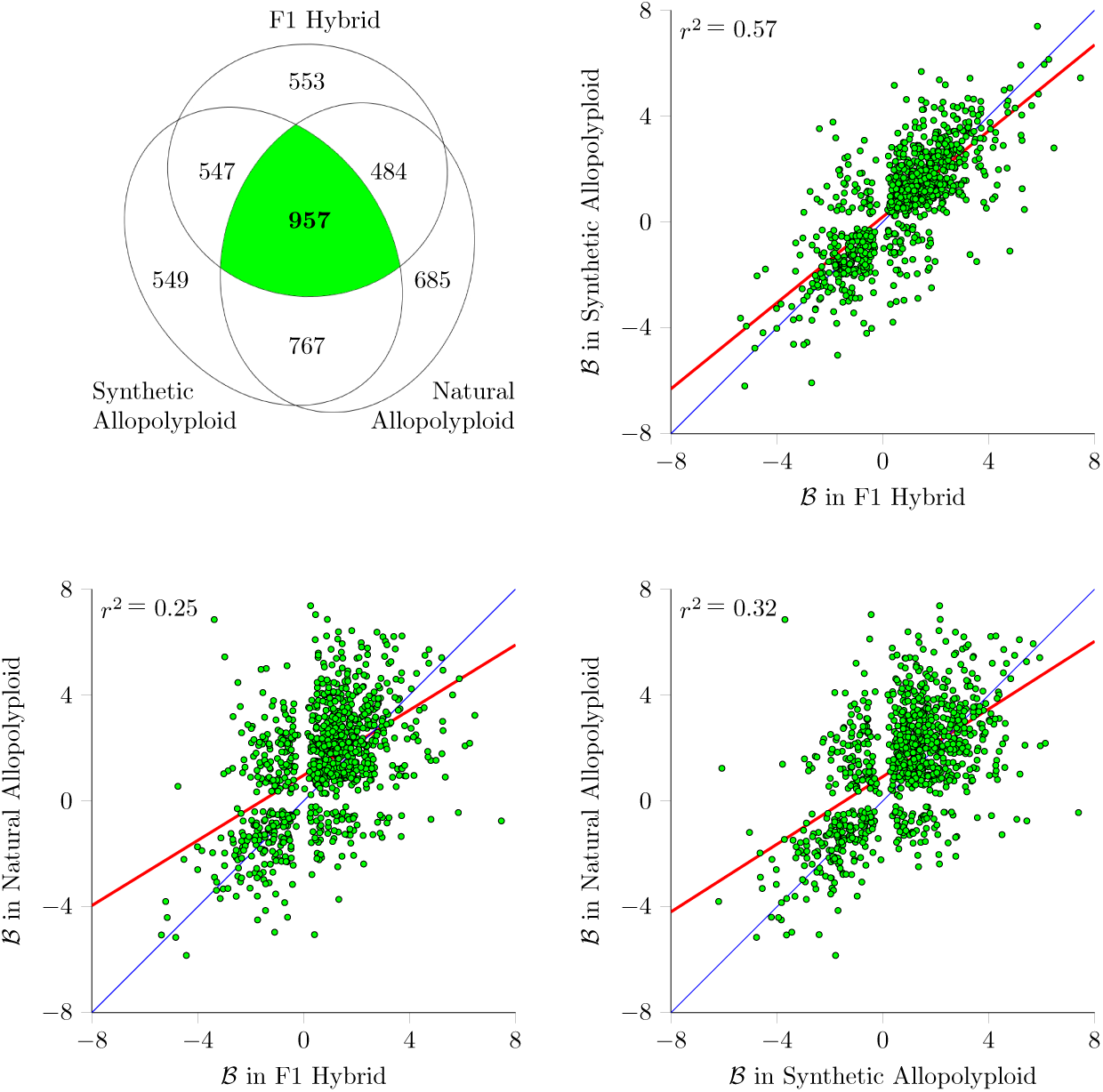
Expression bias in three separate hybrid lineages. (A) Venn diagram of the number of biased homeolog pairs across hybrid lineages (957 homeolog pairs were biased in all three lineages). (B-D) Scatter plots of expression bias (𝓑) for these 957 homeolog pairs comparing hybrid to synthetic allopolyploid, hybrid to natural allopolyploid, and synthetic to natural allopolyploid (red line is linear regression; thin blue line is identity).

#### Transposon density linked to gene expression

One possibility we considered was whether proximal transposon (TE) loads were related to homeolog expression bias. In order to test this it was necessary to annotate TE in *M. guttatus* and *M. luteus* genome assemblies. Using a homology and structured based annotation, as well as *de novo* annotation, we identified the transposons in the *M. guttatus* and *M. luteus* genomes (SI Appendix, Text S1.6). Our survey revealed that 50% of the *M. guttatus* genome assembly is composed of TE sequences that are classified into 863 families. We have compiled a TE exemplar library with 1439 sequences representing the TE composition of the genome (SI Appendix, Text S1.6, Fig. S4). After annotating TEs, in 10kb windows (10kb upstream and downstream of genes as well as within genic region) we calculated the total number of TEs and the number of TE bases. On average, *M. luteus* homeologs and *M. guttatus* homeologs have TE densities of 0.31 and 0.34 (fraction of bases that within a transposon), respectively. In the parents, hybrid, and allopolyploid individuals, this measure of proximal TE density is negatively correlated with gene expression (Fig. 6 and Fig. S5 and S6).

**Figure 6:**
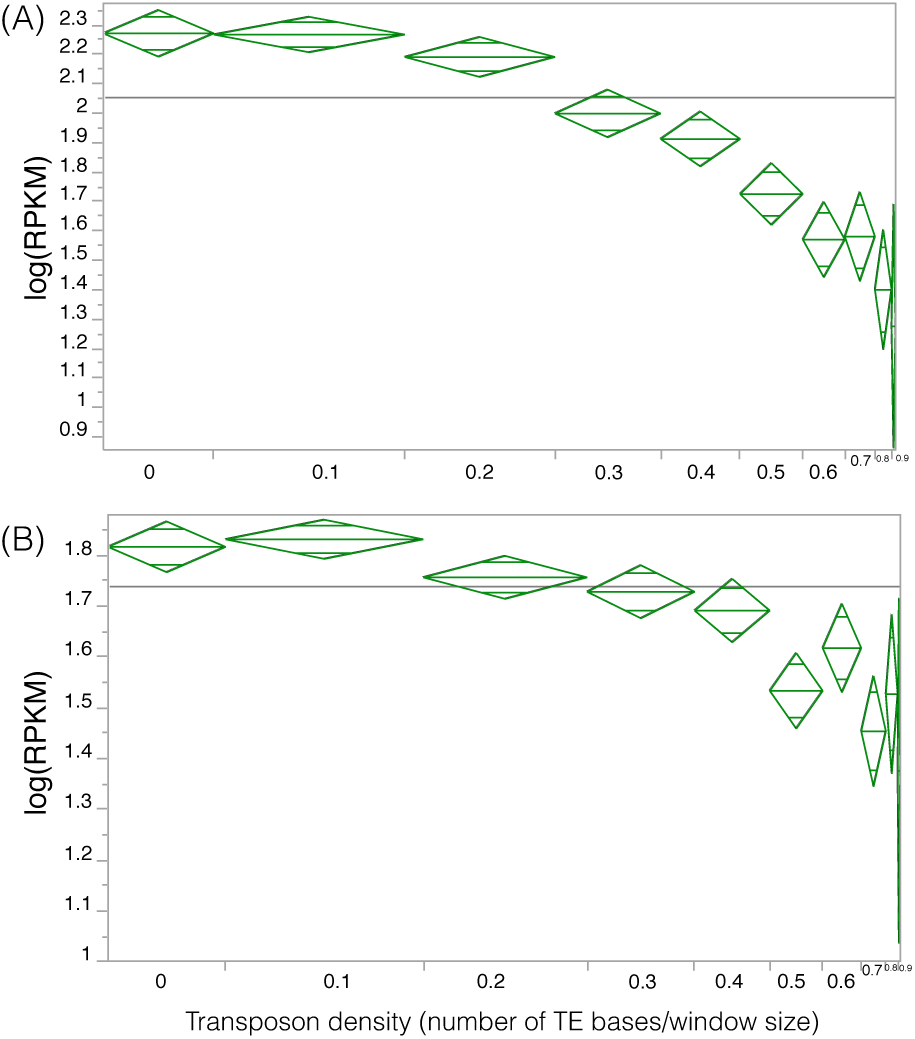
TE density in a window spanning 10kb up and downstream of a gene is negatively related to gene expression. The vertical axis is gene expression in RPKM. The horizontal axis is transposon density, binned into ten windows with width proportional to the number of data points it contains. TE density is negatively related to gene expression in (A) *M. guttatus* and (B) *M. luteus.*

#### Altered DNA methylation patterns in parents, hybrid, and allopolyploids

Building on the finding of expression bias in the hybrid and neo-allopolyploids, we asked whether DNA methylation changes mirror the observed expression bias. First, whole genome bisulfite sequencing (WGBS) was used to determine the methylation status and patterns of methylation change in hybrid and neo-allopolyploid lineages as well as in each parent at CHH (where H = C, A, T), CHG, and CG sites (SI Appendix, Text S1.4). Next, we tested the hypothesis that changes in TE methylation between parents and hybrid or allopolyploids may explain patterns of subgenome dominance.

Methylation patterns in TE and genes and their upstream and downstream regions were compared. CG and CHG methylation patterns in genes and TE are unchanged in the hybrid and allopolyploid lineages (Fig. 7). CHG methylation levels are marginally lower in upstream and downstream regions of genes in the hybrid and slightly higher in the synthetic and natural allopolyploid. CHH methylation levels are decreased signifiantly in upstream and downstream genic regions in the first generation hybrid, decreased slightly in resynthesized allopolyploid, and returned to parental levels in the natural allopolyploid. Similar to findings in genes, transposon bodies and up and downstream regions of transposons are depleted in CHH methylation in the hybrid and synthetic allopolyploid (Fig. 7). Transposon CHH methylation levels are lowest in the first generation hybrid and next highest in the resynthesized allopolyploid. In the natural allopolyploid, CHH methylation of TEs across the *M. guttatus* subgenome returned to near parental levels while *M. luteus* subgenome TE methylation remained lower compared to parental levels. Globally methylation repatterning closely reflects the pattern of homeolog expression bias observed between the two subgenomes in the hybrid and allopolyploids.

**Figure 7:**
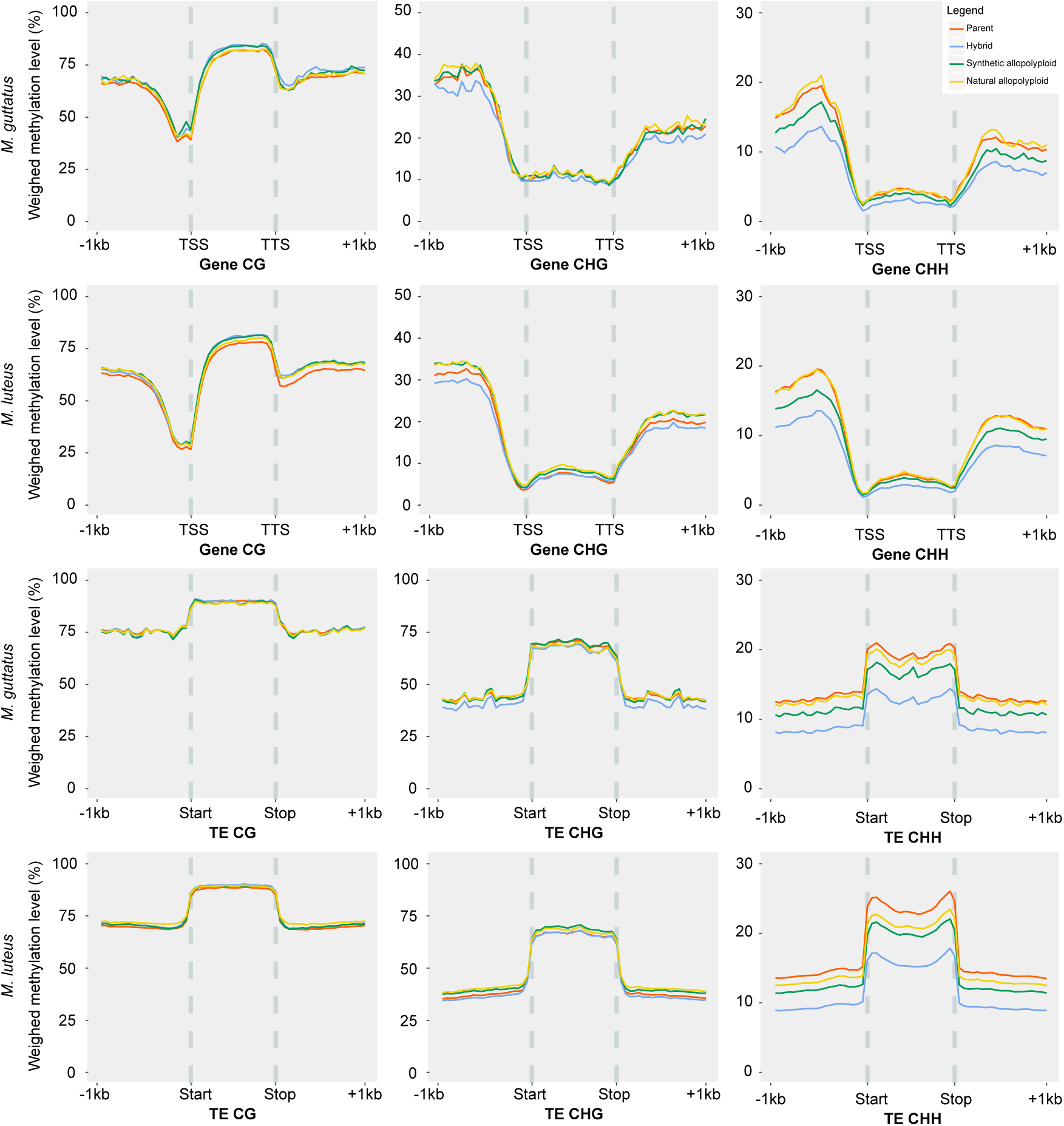
Subgenome specific methylation repatterning in hybrid and allopolyploid *Mimulus.* *M. guttatus* and *M. luteus* subgenome specific patterns of gene (top two rows) and transposon (bottom two rows) methylation. The y-axis is the weighted methylation level. The x-axis shows the gene body (TSS = transcription start site and TTS = transcription termination site) or TE body and one kilobase upstream and downstream. CG, CHG, and CHH methylation levels are shown in the first, second, and third column, respectively. Methylation levels of each individual are shown in unique colors (parents = red; F1 hybrid = light blue; synthetic allopolyploid = dark green; Natural allopolyploid = yellow).

### 3 Discussion

Investigating the aftermath of WGDs across both deep and recent time scales provides clearer insight into the collective evolutionary processes that occur in a polyploid nucleus^46^, including the emergence and establishment of subgenome dominance. Subgenome dominance has largely been investigated in ancient polyploids, including *Arabidopsis*^47^, Maize^34^ and *Brassica*^35^, which revealed the presence of a dominant subgenome with significantly greater gene content and which contributes more to the global transcriptome than the other subgenome(s). Gene expression bias towards one of the subgenomes has also been observed in more recent allopolyploids including those formed as a product of domestication over the past ten thousand years (e.g. wheat^36^ and cotton^26^) and within recently formed natural allopolyploids, namely *Tragapogon mirus*^24^. Due to the recent time scale of these WGD events, gene fractionation bias towards the submissive (non-dominant) subgenomeis not observed. It remains largely unknown how quickly subgenome dominance is established following an allopoly-loid event.

Here we report that subgenome dominance becomes established in the first generation hybrid of two *Mimulus* species. Our analyses show that homeologs from *M. luteus*, compared to *M. guttatus*, are significantly more expressed in the interspecific F1 hybrid and that this expression bias increases over subsequent generations, with the greatest bias observed in the natural (≈ 140 year old) neo-allopolyploid *M. peregrinus.* Using the LRT we determined that the number of biased homeolog pairs also increases with additional generations.

Genome-wide methylation analyses uncovered that CHH methylation levels are greatly reduced in the F1 hybrid. The greatest changes in CHH methylation are observed near gene bodies, near TE bodies, and within TE bodies. The methylation status of many of these CHH sites are regained in the second generation resynthesized allopolyploid; indicating the onset of repatterning of DNA methylation. The methylation status of CHH sites near genes returned to parental levels across both subgenomes in the natural allopolyploid, with regions near *M. guttatus* homeologs being slightly more hypermethylated. Similarly, the CHH methylation status of TEs across *M. guttatus* subgenome returned to near parental levels in the natural allopolyploid. However, this pattern for CHH methylation is not observed across the dominant *M. luteus* subgenome in the natural allopolyloid, with methylation levels within and near TEs remaining noticeably below parental levels. The methylation of CHG sites near TE and gene bodies were also impacted upon hybridization, but to a lesser degree than CHH methylation, and quickly returned to either at or above parental levels in the resynthesized allopolyploid. These observations, a dominantly expressed subgenome with lower TE abundance and lower CHH methylation levels near genes compared to the submissive subgenome, support predictions made by Freeling et al.^41^ to explain subgenome dominance. It is important to note that although *M. luteus* has 84% percent more genes than *M. guttatus*, it only has 41% percent more TEs. This means that the *M. luteus* genome has evolved to have fewer TEs at a genome-wide level.

Our analyses confirm that the density of TEs negatively impacts the expression of nearby genes in *Mimulus*, similar to observations made in *Arabidopsis^40^.* Furthermore, here we show that the repatterning of TE methylation levels are different for the two subgenomes in *M. peregrinus*, mirroring the expression bias observed over the generations following the hybridization. Our results suggest that subgenome dominance may be at least partially due to subgenome-specific differences in the epigenetic silencing of TEs, which was established long ago in the ancestors of the diploid progenitors. The strong correlation between 𝓑 in the F1 hybrid, resynthesized allopolyploid, and independently established natural allopolyploid indicate that the observed subgenome expression dominance is biologically meaningful and likely heritable. The fact that 𝓑 in *Mimulus* hybrids mirrors methylation repatterning and subgenome specific TE densities supports the original hypothesis for the genomic basis of subgenome dominance.

The observed methylation differences between homeologs present on the different subgenomes may represent early earmarks for the ultimate loss (i.e. fractionation) of a duplicate gene copy. Submissive subgenomes in ancient polyploids are more highly fractionated and contribute less to the overall transcriptome compared to the dominant subgenome. Although duplicate genes on either subgenome are not physically lost yet in *M. pereginus*, many homeologs on the submissive subgenome are already functionally absent (low to no expression). Due to selection acting on maintaining proper stoichiometry in dosage-sensitive macromolecular complexes and gene-interaction networks^48 49^, stoichiometric balance is likely best maintained by retaining the more highly expressed copy of interacting genes. One of the biggest opportunities arising from any gene duplication is the possibility of sub- or neo-functionalization. The finding of strong and immediate homeolog expression bias in a hybrid and neo-allopolyploid may have important implications for our understanding of these processes.

In conclusion, there appears to be clear tradeoff between the benefits of epigenetic silencing of TEs (this inhibits their proliferation across the genome) and the effects of TE methylation on neighboring gene expression^40^. Our results support the idea that subgenome dominance may be the result of lineage-specific genomic evolution shaping TE densities and methylation levels. In addition, subgenome expression dominance should not be unique to interspecfic hybrids, but should also occur in intraspecies crosses between lines with different TE loads. These results have major implications to a number of research fields ranging from ecological studies to crop breeding efforts.

### 4 Methods

See supplement for Methods.

